# Attention, Musicality, and Familiarity Shape Cortical Speech Tracking at the Musical Cocktail Party

**DOI:** 10.1101/2023.10.28.562773

**Authors:** Jane A. Brown, Gavin M. Bidelman

## Abstract

The “cocktail party problem” challenges our ability to understand speech in noisy environments, which often include background music. Here, we explored the role of background music in speech-in-noise listening. Participants listened to an audiobook in familiar and unfamiliar music while tracking keywords in either speech or song lyrics. We used EEG to measure neural tracking of the audiobook. When speech was masked by music, the modeled peak latency at 50 ms (P1_TRF_) was prolonged compared to unmasked. Additionally, P1_TRF_ amplitude was larger in unfamiliar background music, suggesting improved speech tracking. We observed prolonged latencies at 100 ms (N1_TRF_) when speech was not the attended stimulus, though only in less musical listeners. Our results suggest early neural representations of speech are enhanced with both attention and concurrent unfamiliar music, indicating familiar music is more distracting. One’s ability to perceptually filter “musical noise” at the cocktail party depends on objective musical abilities.

## 1. Introduction

Background music is a major part of our everyday listening experiences. Listening to music affects our in-store and online shopping behaviors (Ding & Lin, 2012; Garlin & Owen, 2006; North et al., 1999), driving performance (Beh & Hirst, 1999; Cassidy & MacDonald, 2009; Wang et al., 2015), and athletic performance (Atkinson et al., 2004; Chtourou et al., 2012). Listening to speech in background music, however, presents challenges due to the “cocktail party” phenomenon (Cherry, 1953; Haykin & Chen, 2005), in which the listener must attend to one source of auditory input while ignoring competing noise. Listeners can do this by separating the auditory scene into streams in order to isolate target from non-target information (Bregman, 1990).

### 1.1. Effects of background music on speech perception

The impact of background music on concurrent speech or related cognitive tasks is somewhat ambiguous. Background music has been shown to increase listening effort (Du et al., 2020) and performance (Perham & Currie, 2014) on reading comprehension tasks, though other studies have shown no detrimental effect of background music on verbal learning (Jäncke & Sandmann, 2010). Similarly, a meta-analysis (Kämpfe et al., 2011) showed no overall impact of background music on adult listeners across several behavioral domains. While this is in part due to the heterogeneity in experiments investigating background music, it is also worth noting there is significant individual variability in performance. Listeners who prefer not to listen to music while studying showed poorer reading comprehension (Etaugh & Ptasnik, 1982) and more susceptibility to tempo changes in background music (Su et al., 2023). Comprehension was impaired with background music in learners with lower working memory capacity (Lehmann & Seufert, 2017). These effects also vary depending on listener personality, particularly when comparing introverts and extroverts (Avila et al., 2012; Furnham & Allass, 1999; Furnham & Strbac, 2002).

Often taken for granted, the style of the music itself may be an important factor in driving performance, particularly in relation to an individual’s music preference. For example, reading comprehension suffers only when listening to non-preferred music (Johansson et al., 2011), while self-selected music increases task focus and lowers reaction time variability (Homann et al., 2023). However, Perham and Sykora (2012) showed *worse* performance on a letter recall task when listening to preferred music. Genre may also be important. Classical instrumental music can facilitate performance on linguistic accuracy tasks (Angel et al., 2010) and enhance the neural N2 response in an oddball task (Caldwell & Riby, 2007), a marker of perceptual novelty and attentional processing. However, Caldwell and Riby (2007) then showed that at P3, only classical musicians, not rock musicians, showed enhanced processing in classical music. Different genres can induce different moods in listeners (Rea et al., 2010), which can modulate performance on cognitive tasks such as spatial reasoning (Husain et al., 2002). Listeners may also have a preferred tapping tempo, regardless of familiarity to the music (Hine et al., 2022). Thus, studies suggest that personal preference to background music has a significant impact on concurrent behavior and perception.

### 1.2. Acoustic Features

Acoustic features account for a wide range of auditory masking effects and therefore may also influence whether background music hinders speech perception. Intelligibility of conversational speech is worse in piano music played in a low octave and at a faster tempo (Ekström & Borg, 2011), consistent with well-known asymmetries in psychoacoustical masking. Similarly, reading comprehension is worse during very high tempo and louder music (Thompson et al., 2011). This is likely due to the arousal-mood hypothesis (Husain et al., 2002; Thompson et al., 2001), where task performance improves when music increases arousal (and thus induces more positive mood) up to a point, but can then oversaturate, creating states of overarousal that impair performance (Unsworth & Robison, 2016; Yerkes & Dodson, 1908). North and Hargreaves (1999) suggested that high-arousal music requires more cognitive resources than less arousing music. Given that the brain is a limited capacity system, more cognitive resources allocated to music listening means there would be fewer resources left to carry out any concurrent tasks (i.e., speech perception). As a result, cognitive task performance should be worse when background music significantly increases listener arousal.

### 1.3. Vocals

Evidence that background music with vocals impairs concurrent linguistic tasks is clearer. This is likely due, in part, to informational masking, where even unattended sounds in the same domain (e.g., speech on speech) can interfere with target recognition due to lexical interference. Indeed, people listen to instrumental background music while studying or reading, but choose vocal music while driving or performing non-linguistic tasks (Kiss & Linnell, 2022). Music with vocals impaired performance on linguistic tasks (Brown & Bidelman, 2022a, 2022b; Crawford & Strapp, 1994; Scharenborg & Larson, 2018) and immediate recall in learning foreign language task (De Groot & Smedinga, 2014). Importantly, this type of informational masking only occurs when the interfering stream is understood by the listener. Brouwer et al. (2021) showed that an English masker impaired speech intelligibility more than “Simlish,” a fictional gibberish language that shares phonemic patterns with English but lacks semantic meaning. Collectively, these studies suggest that the linguistic status of the background music and degree to which it carries lexical cues can modulate concurrent speech recognition.

### 1.4. Familiarity

Evidence for the role of familiarity in background music is also mixed. Several studies report better performance on language and speech tasks in the presence of familiar compared to unfamiliar background music (Brown & Bidelman, 2022a; Feng & Bidelman, 2015; Russo & Pichora-Fuller, 2008). Presumably, if the listener knows the music, the sequence of the song is predictable, which aids in auditory streaming (Bendixen, 2014). Similarly, if the listener already has mental representations of the familiar music, fewer cognitive resources are needed to process the masking stream and listeners can more easily “tune it out” to prevent interference (Russo & Pichora-Fuller, 2008). Indeed, stream segregation may be easier when the attended and/or the unattended stimuli are more predictable (reviewed in Alain & Arnott, 2000; Jones et al., 1981; Shi & Law, 2010).

However, other studies report more detrimental effects of *familiar* music (Brown & Bidelman, 2022b; De Groot & Smedinga, 2014). Such effects are difficult to explain under the aforementioned arousal hypothesis (Husain et al., 2002) and instead may reflect the redirecting of limited cognitive resources and/or attentional mechanisms (e.g., Lavie, 2005). Familiar music can also provoke autobiographical memories (Belfi et al., 2016; Castro et al., 2020; Janata et al., 2007) and evoke musical imagery (Halpern & Zatorre, 1999; Zatorre & Halpern, 2005), siphoning cognitive resources away from the primary task, and ultimately impairing performance. In support of this notion, we recently demonstrated that speech intelligibility is worse when concurrent background music was familiar to the listener, regardless of whether it contained vocals (Brown & Bidelman, 2022b). The further impairment from vocal music was expected due to informational/linguistic masking.

### 1.5. Musicianship and speech-in-noise (SIN) processing

Another important factor shown to impact cocktail party and SIN listening is musicianship (Bidelman & Yoo, 2020; Yoo & Bidelman, 2019). Several studies report a so-called “musician advantage” in cognitive processing (c.f. Escobar et al., 2019; Hennessy et al., 2022), whereby individuals with musical training show enhancements across domains like audiovisual integration (Wang et al., 2022) and working memory (Brandler & Rammsayer, 2003; Hansen et al., 2013). Musicians are reported to have enhanced auditory skills (Kraus & Chandrasekaran, 2010) supported by a myriad of neuroplastic changes stemming from the cochlea (Bidelman et al., 2016; Bidelman et al., 2017) to cortex (Anderson et al., 2011; Schneider et al., 2002). Musicians are also better at decoding emotion based on speech prosody (Thompson et al., 2004) and have more robust brainstem responses to speech and musical sounds (e.g., Bidelman, Gandour, et al., 2011; Bidelman, Krishnan, et al., 2011; Musacchia et al., 2007). Among the more widely reported—and controversial—musician advantages is enhanced SIN listening (Coffey et al., 2017; Hennessy et al., 2022; Madsen et al., 2017). Musicians are more successful in speech segregation in a multi-talker scene (Baskent & Gaudrain, 2016; Bidelman & Yoo, 2020) and show more resilient subcortical encoding of speech sounds in background noise than nonmusicians (Bidelman & Krishnan, 2010; Parbery-Clark et al., 2009). Listeners with music training are also better able to harness executive control in facilitating auditory attention in SIN listening (Strait & Kraus, 2011), and they are less susceptible to interference from informational masking (Oxenham et al., 2003; Swaminathan et al., 2015).

Importantly, enhanced auditory skills and SIN advantages can be observed in listeners with minimal or no musical training but high levels of innate musicality (e.g., Mankel & Bidelman, 2018; Zhu et al., 2021). This suggests putative benefits in cocktail party listening reported among musicians might not be due to musical training/experience, *per se*, but rather inherent listening skills. Mankel and Bidelman (2018) showed that nonmusicians who scored high on objective measures of musicality had more resilient neural encoding of speech-in-noise than less musical listeners. Similarly, listeners with lower musicality are more affected by the presence of vocals during a speech comprehension task (Brown & Bidelman, 2022a), but only when background music is unfamiliar to them. In contrast, high musicality listeners show less susceptibility to this informational masking effect, indicating that they are more resilient in difficult listening conditions.

### 1.6. Selective attention in cocktail party speech perception

Successful “cocktail party” listening requires successful selective attention (Oberfeld & Klöckner-Nowotny, 2016). Attention also plays a role in auditory stream segregation (Bregman, 1990), although there is some debate whether these streams are created pre-attentively or as a result of attention (Fritz et al., 2007). Such attentional modulation is reflected in the brain as increased activity in auditory cortical areas (Elhilali et al., 2009) with a leftward hemispheric lateralization in cases of speech stimuli (Hugdahl et al., 2003). Neural tracking of the target speech signal is stronger for attended sounds, but the brain still maintains representations of the unattended/background sounds whether they are speech or music (Alain & Woods, 1993; Ding & Simon, 2012; Maidhof & Koelsch, 2011).

Attentional effects can be observed even in the early auditory cortical potentials (ERPs) at sensory stages of speech processing. There is a long-established attentional enhancement of the auditory N1, a negative peak around 100 ms in the canonical auditory ERP (Ding & Simon, 2012; Hillyard et al., 1973; Woldorff et al., 1993). However, attentional modulation of cortical responses has also been observed as early as 40 ms (Teder et al., 1993; Woldorff et al., 1993; Woldorff & Hillyard, 1991) and 75 ms (Bidet-Caulet et al., 2007). These findings suggest attention exerts early influences on auditory sensory coding which may improve SIN analysis by bolstering and/or attenuating target from non-target streams in a cocktail party scenario.

In a study by Ding and Simon (2012), listeners were instructed to attend to one of two speakers. Neural representations of both the attended and unattended talkers were preserved but heavily modulated by attention; that is, cortical encoding of the attended speaker was much larger. Our previous study (Brown & Bidelman, 2022a) similarly measured neural tracking to continuous speech, but that study manipulated attention by changing the background music familiarity. The current study extends these results by forcing listeners’ attention (as in Ding & Simon, 2012) to measure similar tracking ability for speech on music rather than often-used speech on speech.

The current experiment aims to elucidate speech perception in background music and how it is modulated by (i) forced attention, (ii) familiarity of the music, and (iii) listeners’ musicality. Participants listened to a speech audiobook and concurrent familiar/unfamiliar music while completing a keyword identification task that forced attention to either the continuous speech or the song lyrics. We measured neural activity using multichannel EEG and extracted the brain’s tracking of the continuous amplitude envelope of the audiobook and song vocals using temporal response function (TRF) analysis. We hypothesized that (1) keyword identification and neural tracking would be worse for speech presented in background music compared to in silence (i.e., expected masking effect); (2) neural speech tracking would be weaker in unfamiliar background music (Brown & Bidelman, 2022a); (3) speech tracking would be enhanced when speech was the attended condition versus music as the attended condition; and (4) less musical listeners would show poorer attentional juggling between the speech and music attention conditions, suggesting worse attentional allocation of cognitive resources.

## 2. Materials and Methods

### 2.1. Participants

The sample included 31 young adults ages 21-33 (*M* = 24, *SD* = 3.3 years, 13 male). All participants showed normal audiometric thresholds < 15dB HL at octave frequencies 250-8000 Hz, as well as normal SIN perception (QuickSIN scores < 3dB SNR loss; Killion et al., 2004). All reported English as their native language. Participants were primarily right-handed (mean 70% laterality using the Edindurgh Handedness Inventory; Oldfield, 1971). Participants also self-reported years of formal music training, which ranged from 0 to 16 years (*M* = 4.9 years, *SD* = 4.92). Each was paid for their time and gave written consent in compliance with a protocol approved by the Institutional Review Board and the University of Memphis.

### 2.2. Stimuli

#### 2.2.1. Music

We used unfamiliar and familiar pop song music selections. To qualify as “familiar,” the song had to appear on the Billboard Hot 100 list (https://www.billboard.com/charts/hot-100/) at least once. Each song was sung by a female singer. All songs were performed at a tempo from 110-130 beats per minute. Thompson et al. (2011) showed an effect of faster musical tempi on concurrent reading comprehension, so the tempo range here falls in the “slow” to “intermediate” range of their experiment to avoid tempo effects. Using the above criteria, four songs were used in the current experiment: two familiar (“*Girls*” by Beyonce; “*Stronger (What Doesn’t Kill You)*” by Kelly Clarkson) and two unfamiliar (“*Joan of Arc on the Dance Floor*” by Aly & AJ; “*OMG What’s Happening*” by Ava Max). Familiarity categories were determined using a pilot study (*N* = 37, 15 males, 22 females; age *M* = 26, *SD* = 2.95). Participants were asked to rate several songs on a 5-point Likert scale from “Not familiar at all” to “Extremely familiar.” The songs used in the current EEG experiment were the two most and least familiar songs from those pilot results.

Songs were converted from stereo to mono, sampled at 44100 Hz, and truncated from onset to 2 min. To maximize data available for analysis, instrumental introductions were cut so that vocals began withing 2 sec of the start of the clip. Clips were RMS-normalized to equate overall levels. However, amplitude fluctuations in the music (i.e., short instrumental segments, chorus) were allowed to vary within 10 dB of the overall RMS to maintain the natural amplitude envelope of the original music.

#### 2.2.2. Speech

The speech stimulus was a public domain audiobook from LibriVox (https://librivox.org/). The selected audiobook was “*The Forgotten Planet*” by Murray Leinster read by a male speaker; importantly, this story was unfamiliar to all participants. The story was separated into 36, 2-min segments. Silences longer than 300 ms were shortened to avoid long gaps in the speech (Brown & Bidelman, 2022a; Ding & Simon, 2012).

### 2.3 Task

During EEG recording (described below), each audiobook story clip was presented concurrently with one of the four songs in a random order at a signal-to-noise ratio (SNR) of 0 dB or in silence. The story clips were presented in sequence but were broken up into 8 blocks to allow breaks during the task. For half the experiment, the participant was instructed to attend to the audiobook and listen for a keyword; they were instructed to quickly press the space bar every time they heard the keyword. The other half of the experiment was identical, but listeners were cued to listen for a keyword in the music song vocals. After completing the experiment, participants indicated their familiarity with each song on a sliding scale from 0 (not familiar) to 10 (extremely familiar). They were also asked how much they liked each song (0 to 10 scale).

Participants also completed the shortened version of the Profile of Musical Perception Skills (PROMS-S; Zentner & Strauss, 2017) to assess music-related listening skills. The PROMS is broken up into several subtests that assess different perceptual functions related to music (e.g., rhythm, tuning, melody recognition). In each subtest, two tokens (e.g., rhythms or tones) are presented, and the listener must judge whether the tokens are the same or different using a five-point Likert scale (1 = “definitely different”, 5 = “definitely same”).

### 2.4 EEG recording and preprocessing

Participants sat in an electrically shielded, sound-attenuated booth for the duration of the experiment. Continuous EEG recordings were obtained from 64-channels with electrode position according to the 10-10 system (Oostenveld & Praamstra, 2001). Neural signals were digitized at a 500 Hz sample rate using SynAmps RT amplifiers (Compumedics Neuroscan, Charlotte, NC, USA). Contact impedances were maintained below 10kΩ. Music and speech stimuli were each presented diotically at 70 dB SPL (0 dB SNR) via E-A-RTone 2A insert headphones (E-A-R Auditory Systems, 3M, St. Paul, MN, USA). Presentation of the speech alone served as a control condition to assess speech tracking without music. Stimuli were presented using a custom MATLAB program (v. 2021a; MathWorks, Natick, MA, USA) and routed through at TDT RP2 signal processor (Tucker-Davis Technologies, Alachua, FL, USA).

EEGs were re-referenced to the average mastoids for analysis. We visually inspected the power spectrum for each participant’s recording via EEGLAB (Delorme & Makeig, 2004) and paroxysmal channels were spline interpolated with the six nearest neighbor electrodes. The cleaned continuous data were then segmented into 2-minute epochs. Data from 0 to 1000 ms after the onset of each epoch were discarded in order to avoid transient onset responses in later analyses (Crosse et al., 2021). Epochs were then concatenated per condition, resulting in 16 min of EEG in each attention condition for each familiarity condition.

### 2.5. Data analysis

#### 2.5.1. Behavioral data analysis

Keypresses were logged and compared to the onset of each keyword. A press that fell within 300-1500 ms after the onset of the word was marked a “hit.” Responses earlier than 300 ms were discarded as improbably fast guesses (e.g., Bidelman & Walker, 2017). A keyword with no response in the window was marked a “miss,” and a response not in a keyword window was marked as a “false alarm.” Hits and false alarms were used to calculate *d’* (d-prime) sensitivity. *d’* was calculated by subtracting the z-score of the false alarm rate from the z-score of the hit rate. Because values of 0 or 1 cannot be z-transformed, hit rates or false alarm rates of 0 were changed to 0.001, and rates of 1 were changed to 0.99 to allow for calculation of *d’* (Macmillan & Creelman, 2005).

#### 2.5.2. Temporal response functions (TRFs)

We quantified the neural tracking to the continuous speech signal using the Temporal Response Function toolbox in MATLAB (Crosse et al., 2016). The TRF is a linear function that models the deconvolved impulse response to a continuous stimulus. We extracted the temporal envelope of the continuous audiobook speech via a Hilbert transform. EEG recording data were down-sampled to 250 Hz, then filtered between 1 and 30 Hz to target cortical activity to the low-frequency speech envelope. EEG and stimulus data were both z-score normalized. Due to inherent inter-subject variability, we computed a TRF for each individual (Crosse et al., 2016). We used 6-fold cross-validation to derive TRFs per familiarity and attention condition, then used ridge regression to find the optimal λ smoothing parameter (Crosse et al., 2021). The model was first trained on the neural response to the attended speech-in-quiet condition to find optimized λ parameter, which was the value that resulted in the maximum reconstruction accuracy. That parameter was then used to compute TRFs for the other masking and attention conditions. This approach avoids overfitting while preserving individual response consistency and increasing decoding accuracy across all speech-tracking conditions (Simon et al., 2023). We trained the model using EEG recordings from a fronto-central electrode cluster (F1, Fz, F2, FC1, FCz, FC2, C1, Cz, C2) to further optimize fit based on the canonical topography of auditory ERPs.

From TRF waveforms, we measured the amplitude and latency of the “P1” and “N1” waves, which occur within the expected timeframe of auditory attentional effects in the ERPs. P1_TRF_ was measured as the positive-going deflection at ∼50 ms and N1_TRF_ as the negative peak around ∼100 ms. We measured RMS amplitude and latency for each peak.

#### 2.5.3. Statistical analysis

Statistics were computed in R using the *lme4* package (v. 1.1.32; Bates et al., 2015). We used mixed models with combinations of familiarity, attention, and PROMS level as fixed effects and subject as a random factor. Effect sizes are reported as partial eta squared computed from the *emmeans* package (v. 1.8.5; Lenth, 2023). Multiple comparisons were adjusted using Tukey corrections.

In preliminary analyses we also examined TRFs at two frontal clusters over the right (Fz, F2, F4, F6, F8, FC6, FT8) and left (Fz, F1, F3, F5, F7, FC5, FT7) scalp to investigate any hemisphere differences. There were no significant interactions between hemisphere and attention for P1_TRF_ (amplitude: *p* = 0.94; latency: *p* = 0.34) or N1_TRF_ (amplitude: *p* = 0.89; latency: *p* = 0.86), so subsequent analyses and figures use the frontal central cluster that was used to train the TRF model.

## 3. Results

### 3.1. PROMS musicality scores

PROMS scores ranged from 24.5 to 59, (*M* = 40.45, *SD* = 9.81) and were significantly positively correlated with listeners’ years of formal music training (*r*(30) = 0.569, *p* < 0.001). As in previous studies (Brown & Bidelman, 2022a; Mankel & Bidelman, 2018), we used a median split to create “high PROMS” and “low PROMS” groups. These groups do not necessarily reflect years of musical training (“musicians” vs. “non-musicians”), but rather, an objective measure of listeners’ musicality (i.e., music perceptual skills).

### 3.2. Masking effect

Figure 1 shows the effect of masking on speech processing. Keyword tracking performance was significantly worse in speech during concurrent music than in quiet (*F*(1,115) = 52.31, *p* < 0.001, η^2^_p_ = 0.31). Paralleling behavior, the neural TRF P1_TRF_ to speech was longer in latency for masked speech than clean speech (*F*(1,115) = 4.78, *p* = 0.003, η^2^_p_= 0.07), indicating poor encoding of the target speech envelope in noise. The findings confirm our masking manipulation was successful in weakening the behavioral and neural representation for speech with music as a background noise.

**Figure 1.**
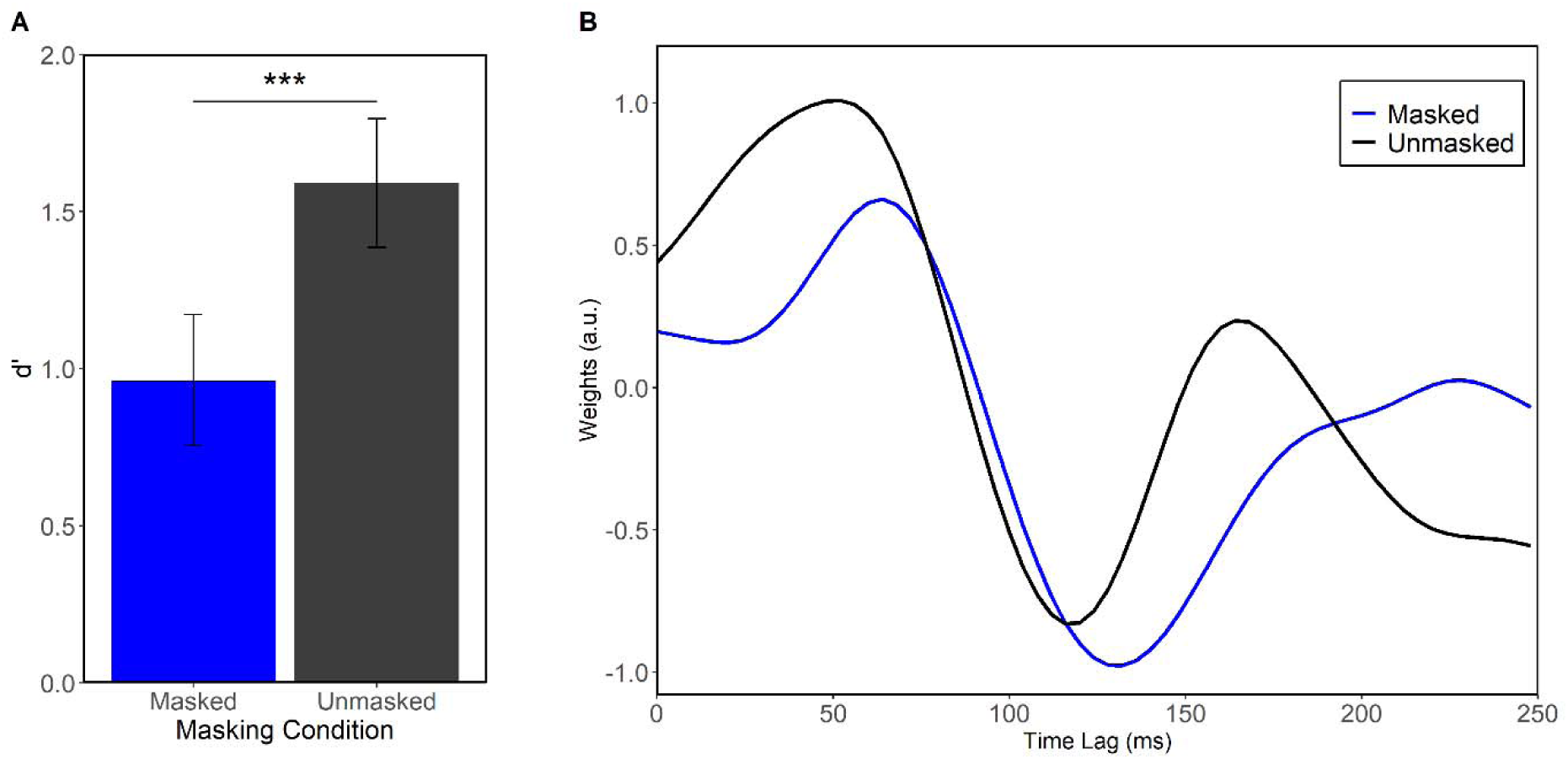

### 3.3. Familiarity effect

When separating the music maskers by familiarity, we found a significant effect of familiarity in the strength of the P1_TRF_ evoked by speech (Figure 2). Amplitude was larger in unfamiliar music than in familiar (*F*(2,137) = 3.21, *p* = 0.043, η^2^_p_= 0.04). There were no amplitude differences between unfamiliar and speech in quiet (*p* = 0.93) or between familiar and quiet (*p* = 0.24). There was also an effect of latency (*F*(2,110) = 4.25, *p* = 0.015, η^2^_p_= 0.07), which reflected the masking effect. Post-hoc Tukey tests showed that latency of the speech-P1_TRF_ in familiar music was longer than speech in quiet (*t*(110) = 2.73, *p* = 0.020). The same prolongation was true for unfamiliar music (*t*(110) = 2.67, *p* = 0.024). There were no differences at N1_TRF_.

**Figure 2.**
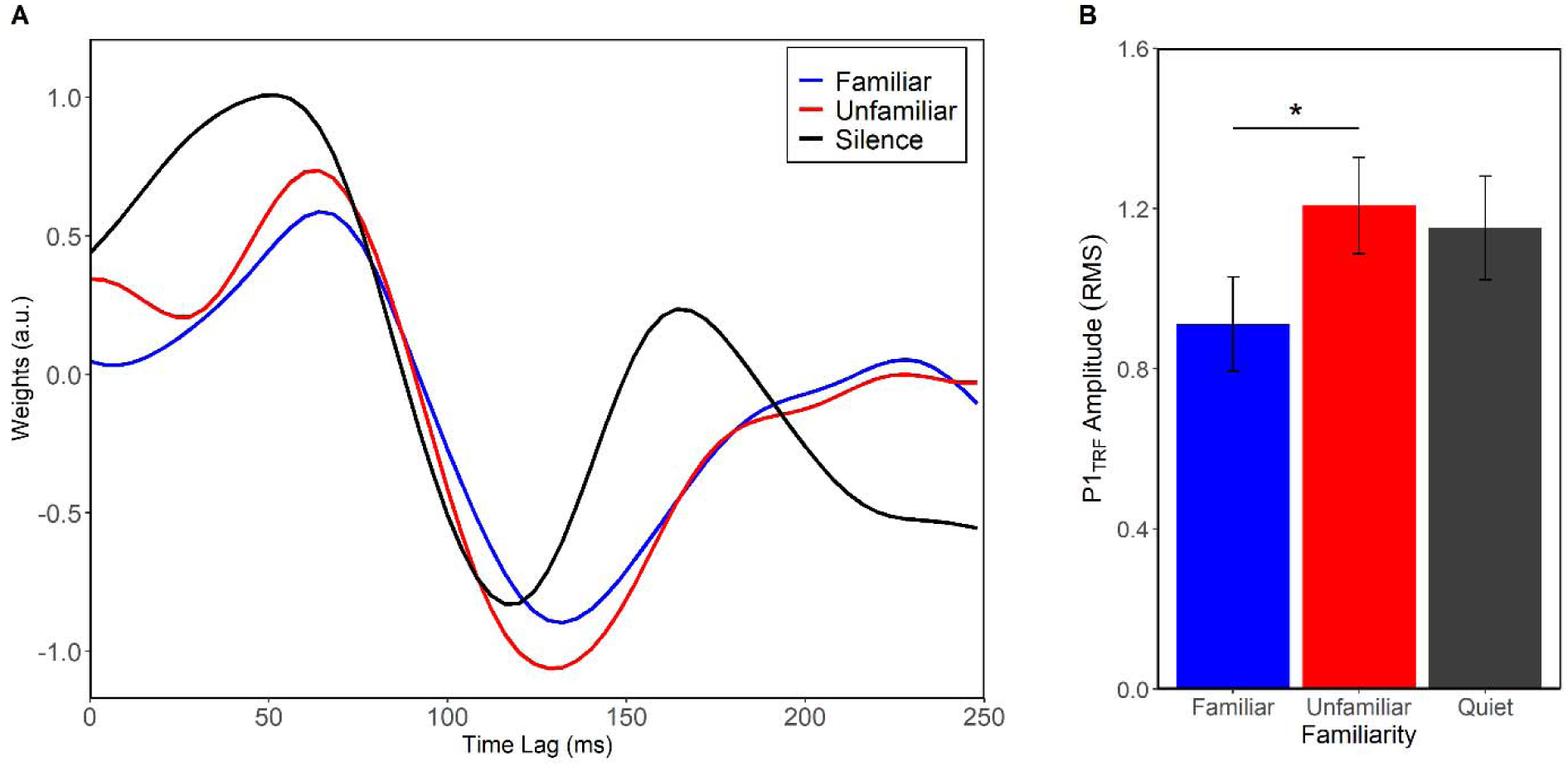
Speech encoding differs between familiar and unfamiliar music maskers. (A) Grand average TRFs (fronto-central electrodes) representing the neural tracking of speech in familiar and unfamiliar background music. (B) P1_TRF_ was larger when presented with unfamiliar music. Error bars represent ± 1 s.e.m. \**p* < 0.05

### 3.4. Attention

We found a significant effect of forced attention on TRF *speech tracking* (Figure 3) dependent on whether listeners were attending to the speech or song vocals. Notably, TRFs were evident in both conditions suggesting the neural representation of continuous speech was maintained whether or not it was the attended stream. However, N1_TRF_ responses were earlier when attending to the speech compared to song (*F*(1,195) = 9.59, *p* = 0.002, η^2^_p_ = 0.05), indicating speech tracking was enhanced by attention. There was no difference in N1_TRF_ amplitude nor latency/amplitude at P1_TRF._

**Figure 3.**
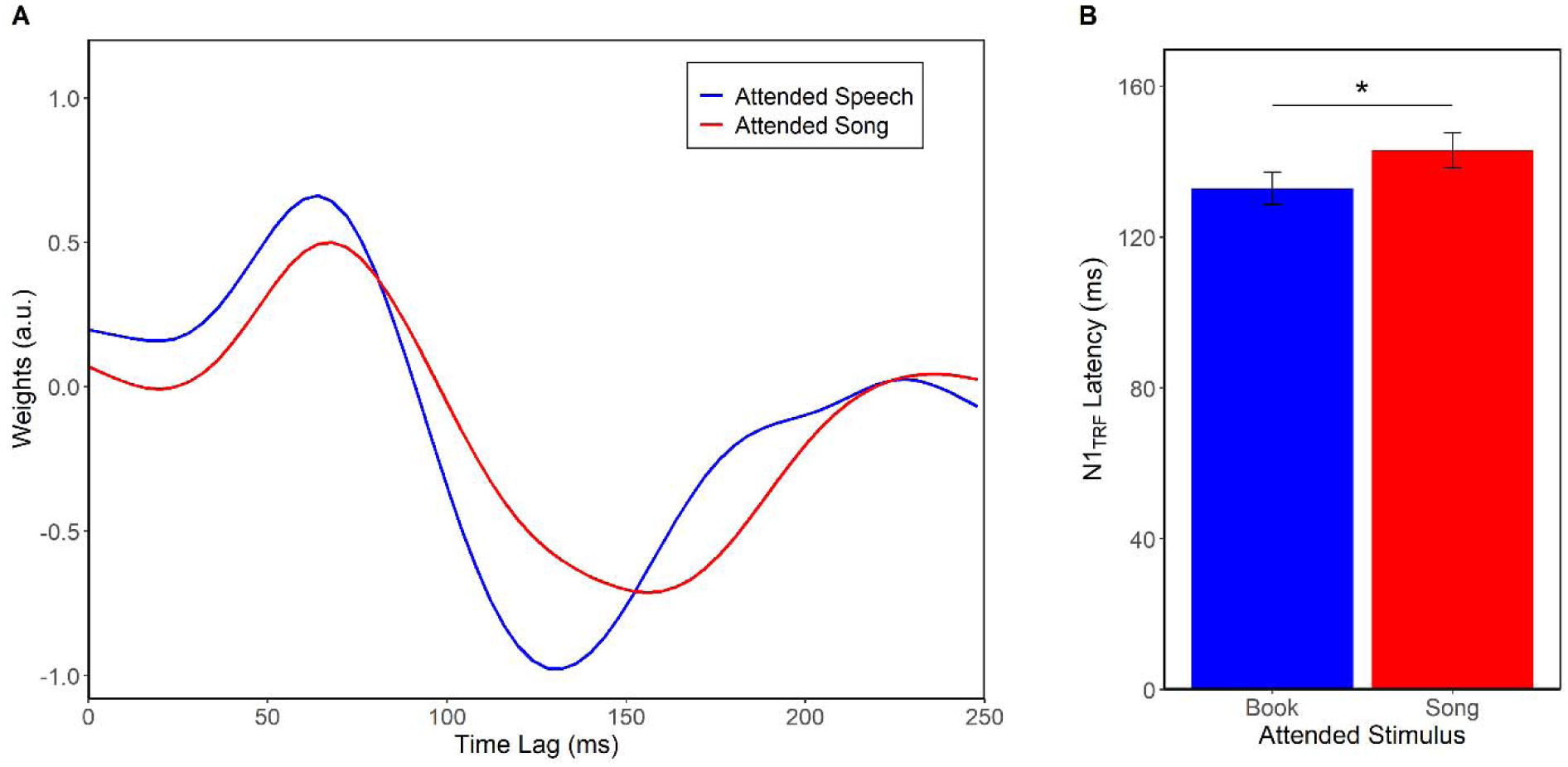
Selective attention modulates neural speech encoding. (A) Grand average TRFs (plotted at fronto-central electrode cluster) for speech tracking when attention is forced to speech versus song. (B) N1_TRF_ for speech encoding is prolonged when attending to the music. Error bars represent ± 1 s.e.m. \*\**p <* 0.01

### 3.5. Effects of musicality

To investigate the relationship between attention and musicality (Figure 4), we split the sample based on a median split of the PROMS musicality scores and examined *a priori* contrasts for the attentional effect in the low PROMS and high PROMS groups. In the low PROMS group, N1_TRF_latency was longer when attending to the song than when attending to speech (F(1,14)= 13.37, *p* = 0.003, η^2^_p_ = 0.49). In stark contrast there was no N1_TRF_ latency difference in the high PROMS group (*p* = 0.42), suggesting the neural tracking of speech was equally good whether or not it was the attended stream.

**Figure 4.**
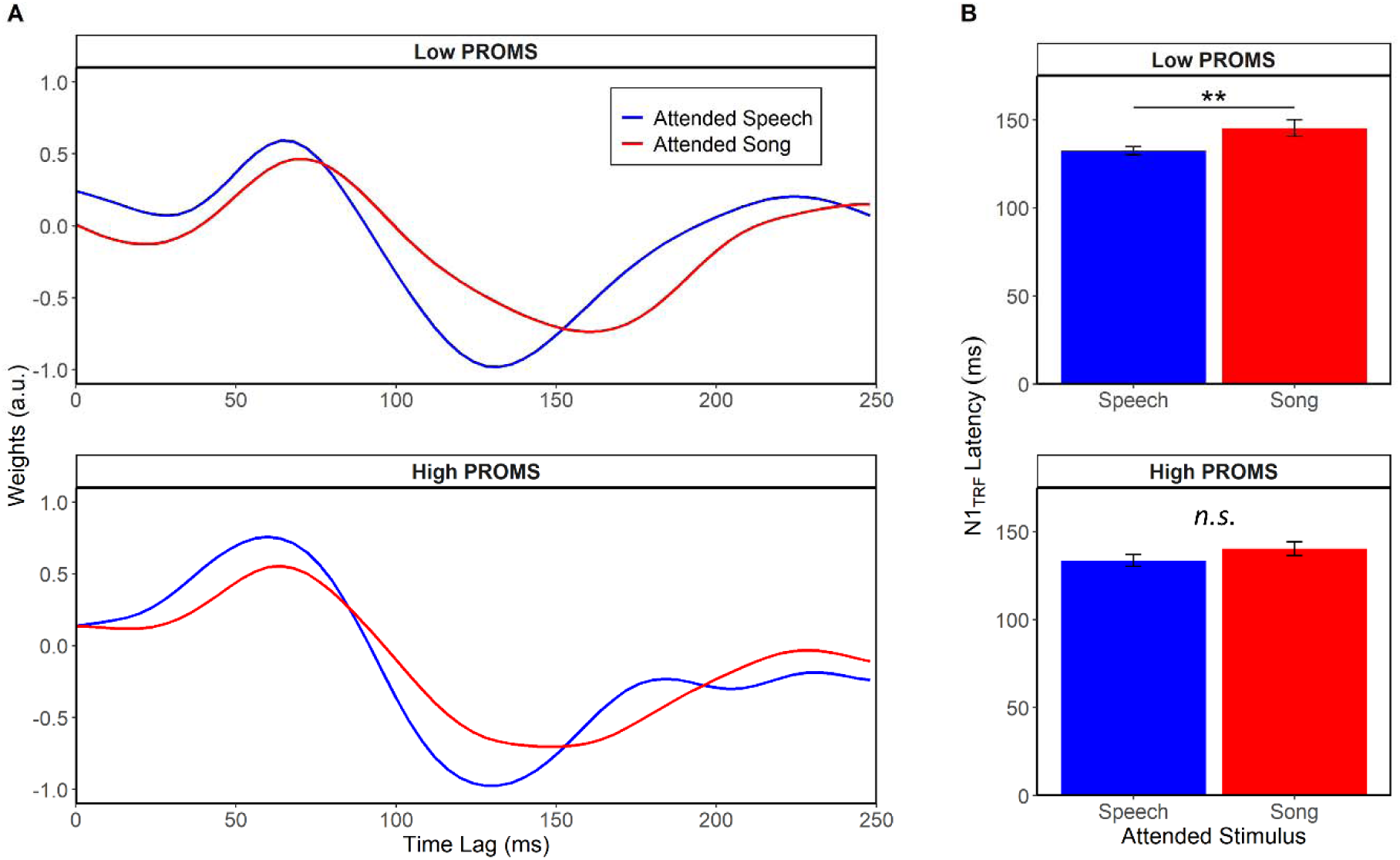
Attentional allocation at the cocktail party differs between less and more musical listeners. (A) TRF waveforms tracking to speech for low vs. high PROMS listeners. (B) N1_TRF_ responses were later than when attending to speech, but only for the less musical listeners. There was no difference in the high PROMS group between music and speech attend conditions. Error bars represent ± 1 s.e.m. \*\**p* < 0.01

When visualizing the N1_TRF_ latency differences between PROMS groups, it was clear the lack of effect in the high PROMS listeners was due to their greater inter-subject variability. To further investigate this, we calculated a “divided listening index” for each listener by taking the latency difference between forced attention to song vocals and forced attention to speech (i.e., N1_song_ - N1_speech_) (Figure 5). Positive values indicate longer latencies when attending to the song vs. attending to speech (i.e., attend song > attend speech; as in Fig. 3A), and thus more susceptibility to music-on-speech masking; negative values indicate longer latencies when attending to speech vs. attending to the song (i.e., attend speech > attend song).

**Figure 5.**
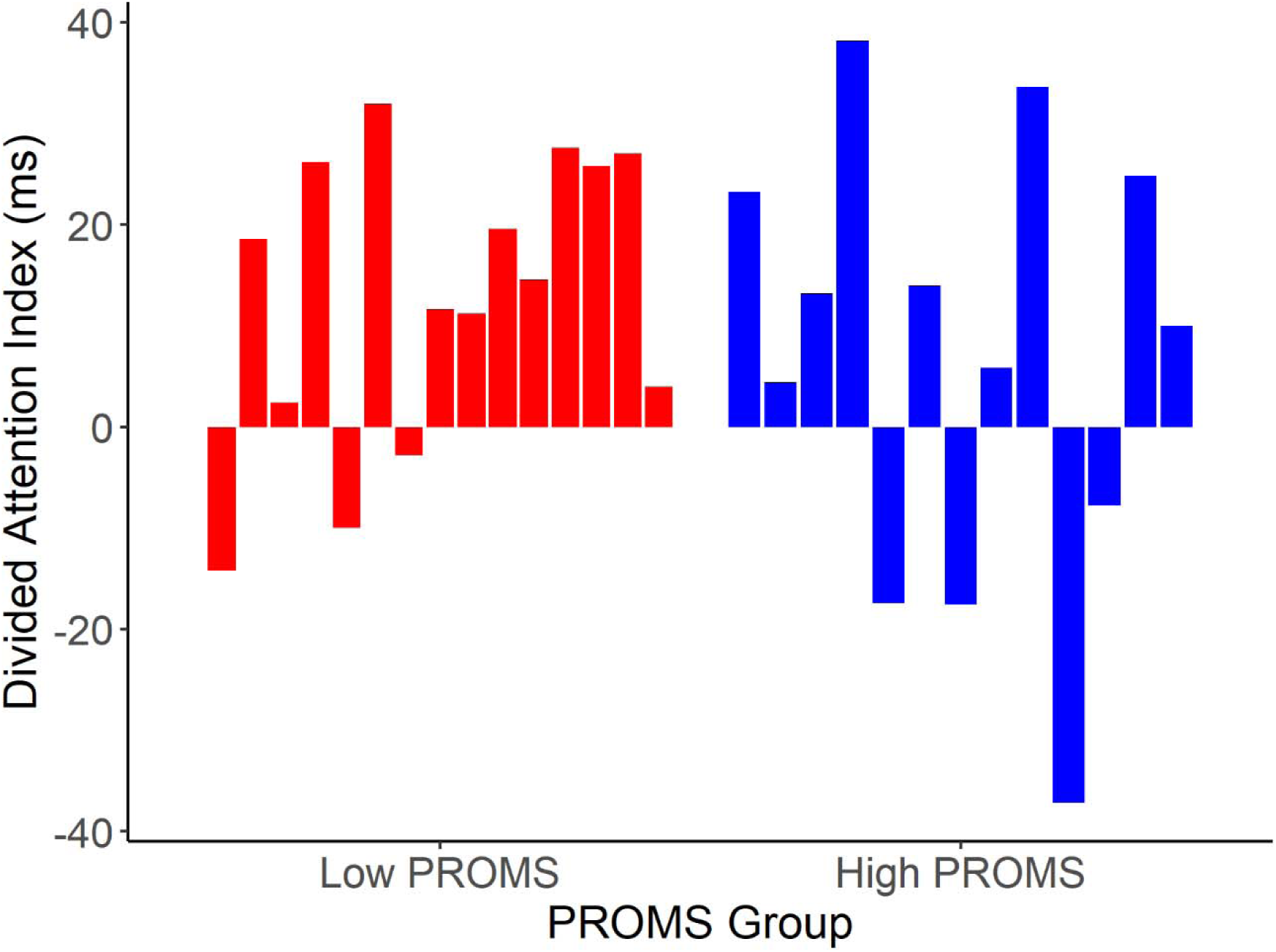
Divided attention index varies in low and high PROMS listeners. Each bar corresponds to one participant and is the difference between speech tracking N1_TRF_ latency when attending to song and attending to speech. A positive index indicates a longer response latency when tracking *speech* when attending to the background *music*.

## 4. Discussion

In this EEG study, participants listened to speech-music cocktail party mixtures (audiobook + pop music) while they selectively attended to either the speech or the song lyrics. We measured neural tracking of the temporal speech envelope of continuous speech using temporal response functions (TRFs). Beyond expected masking effects of concurrent music, we found early cortical responses (P1_TRF_; ∼ 50 ms) to attended speech were larger when the background music was unfamiliar to the listener. Neural responses also showed strong attentional effects, where N1_TRF_ (∼100 ms) to speech was later when attending to song than attending to speech in speech-music mixtures. Interestingly, this attention difference was only prominent in less musical listeners; more musical listeners showed more resilience in tracking speech regardless of whether it was the attended or non-attended stream. Our findings highlight that parsing speech at the cocktail party depends on both the nature of the music noise backdrop itself as well as the perceptual expertise of the listener.

### 4.1. Attention enhances neural speech tracking in musical noise

We found a prolonged N1_TRF_ for speech tracking when the audiobook is the attended stream rather than the background (i.e., when attending to the song lyrics). Our far-field EEG data agree with intracranial recordings which show spectrotemporal representations of speech in auditory cortex are heavily modulated by attention (Mesgarani & Chang, 2012). Using spectrotemporal response functions (STRF) applied to far-field MEG, Ding and Simon (2012) showed similar attention effects at 100 ms (M100_STRF_) in a two-talker selective attention task where responses were stronger for the attended speaker versus the unattended speaker. Our similar findings at comparable effect sizes (present study: η^2^_p_ = 0.05; Ding and Simon (2012): η^2^_p_= 0.06) show that these attention effects replicate across domains (speech/speech versus speech/music).

It is clear that some representation of the ignored stimulus is created and is weaker than that of the attended (Brodbeck et al., 2020; Ding & Simon, 2012), but to what extent, or by what mechanism, is still unclear (Zion-Golumbic & Schroeder, 2012). While there is an unattended stream representation, it may not be as processed as the target stream (Cusack et al., 2004). Attention creates a hierarchy of processing where only attended streams are fully segregated and elaborated. Background music may not be segregated into different streams (i.e., different musical instruments may not form different streams). Multivariate TRFs (Crosse et al., 2021) to several acoustic or musical features could help to assess relevant salience of those features and give insight into how the unattended music is parsed.

Speech intelligibility is easier when the target and interfering speakers are spoken by different-sex speakers due to differences in voice fundamental frequency (Brungart, 2001). Bregman (1990) made the distinction between segregation (differentiating different targets or talkers) and streaming (continuously tracking the separated elements). The current study focused on continuous streaming, so segregation was facilitated by having different-sex stimuli (female vocalists, male audiobook reader). The aim of this experiment was not to look at acoustic differences in segregation, but in attentional streaming effects. Future studies may use same-sex stimuli (e.g., a male speaker and a male vocalist) to further investigate speech/music stream segregation when the target and maskers are more similar.

### 4.2. Early cortical speech processing is weaker in familiar music

We found that P1_TRF_ to continuous speech was smaller when presented with familiar music. Previous studies from our lab (Brown & Bidelman, 2022a, 2022b) have investigated the role of familiarity in background music on concurrent speech perception using ecological music stimuli like those here. Both studies identified speech processing differences between familiar and unfamiliar music maskers. We previously reasoned that those differences were the result of different allocations of limited cognitive resources needed to facilitate selective attention and inhibit the music maskers (Kahneman, 1973; Lavie et al., 2004). However, prior studies did not force attention to speech and music (only speech was tracked behaviorally), so such explanations were only speculative. Our data here confirm the impact of background music on speech processing most probably results from subtle changes in the spotlight of attention as familiar music draws attention away from the primary speech signal. These findings agree with other work showing neural synchronization is stronger for familiar than unfamiliar music (Weineck et al., 2022). Stronger synchronization to familiar music would tend to reduce entrainment to other concurrent signal, as observed here for speech.

While we favor explanations based on attention, familiarity effects could instead result from idiosyncratic acoustic differences between music selections. However, we aimed to combat this by using multiple songs per familiarity condition, as well as using several criteria to match the different songs: genre, tempo, gender of vocalist, key, and beat strength (i.e., pulse; Lartillot et al., 2008). Additionally, we have investigated the role of several acoustic factors, including pulse, on similar familiarity findings and found that while there were acoustic drivers of those effects, the effect sizes were several orders of magnitude smaller than those of music familiarity (Brown & Bidelman, 2022b). Future studies using this paradigm could use multivariate TRFs (Crosse et al., 2016) to see which acoustic variables contribute more to perceptual tracking (e.g., amplitude envelope to vocals and to full song, spectral flux of full song, etc.).

The early P1 effects in our data contrast several MEG studies that have not shown attentional modulation in auditory cortical processing before 100 ms (Akram et al., 2017; Chait et al., 2010; Ding & Simon, 2012; Fujiwara et al., 1998; Miran et al., 2018; Puvvada & Simon, 2017). Several explanations may account for differences between this and previous studies. First, the P1 component at 50 ms is thought to be generated by lateral superior temporal gyrus (Liégeois-Chauvel et al., 1994; Ponton et al., 2000) with radial oriented current dipole. MEG is relatively insensitive to radial currents (Scherg et al., 2019), which might explain why MEG TRF studies have not observed attentional modulation in the P1. Second, P1 is a small amplitude component of the auditory ERPs that is quite variable at the single-subject level. The earlier familiarity effects observed in this (P1_TRF_) compared to previous work (N1_TRF_) could be due to the larger sample of the current study. Nevertheless, the presence of familiarly-attention effects at ∼50 ms suggests music (and how familiar it is to the listener) exerts an influence on speech coding no later than primary auditory cortex (Picton et al., 1999).

Interestingly, Yang et al. (2016) showed that musicians’ performance on cognitive tasks was worse when the background music was played on their trained instrument (e.g., a trained pianist performed more poorly on a verbal fluency test when the background music was played on a piano versus a guitar). If we assume their chosen instrument is more “familiar” to them, then these findings contrast our data. In our previous study (Brown & Bidelman, 2022a), we found more musical listeners were less impacted by background music familiarity. Here, familiarity was measured by self-report and presumably based on real-world exposure to the songs. The operational definition of “familiar” ranges across studies, from real-life exposure (Russo & Pichora-Fuller, 2008) to in-lab training (Weiss et al., 2016) to real vs. artificial instrument timbre (Van Hedger et al., 2022). Further research in this area should carefully consider that definition.

### 4.3. Musicality impacts attentional allocation

The N1_TRF_ peak in response to speech was prolonged when attention was directed to the song versus towards the speech. However, we only see this difference in the low PROMS group (i.e., the less musical listeners), which is likely due to the variance in the high PROMS group (i.e., the more musical individuals). The variability in divided attention index for the high PROMS group may indicate possible differences in listening strategies as a function of listeners’ musicality and/or specific instrument of training. Indeed, several of the high PROMS listeners who showed a negative divided listening index (i.e., speech-tracking latency was longer when it was *not* the attended condition) reported training in non-ensemble instruments, while those with a positive index tended to be ensemble instrumentalists. In general, high musicality listeners show less change between attend speech and attend music conditions, indicating they were more successful in tracking speech regardless of whether or not it was in the attentional spotlight. Similarly, the larger attention-dependent change in TRFs of low PROMS listeners suggests they are more susceptible to changes in background music, possibly resulting from poorer attentional resource allocation and/or increased distractibility by the background (Brown and Bidelman, 2022a). The forced attention manipulations in the current study create new evidence for this explanation. Here, low PROMS listeners showed worse inhibition of the background music suggesting less musical listeners are poorer at regulating auditory attention. In this vein, attentional benefits are observed in trained musicians (Strait et al., 2010; Thompson et al., 2017; Yoo & Bidelman, 2019) and improvements in selective attention might also account for individual differences in cocktail party listening (Oberfeld & Klöckner-Nowotny, 2016). Musical training also correlates with better tracking of the to-be-ignored stream, as well as a more balanced representation of the attended and to-be-ignored streams (Puschmann et al., 2019). These studies, along with current data, support the link between musicality, attentional deployment, and cocktail party listening.

Collectively, our PROMS group differences imply that listeners might approach the speech-music cocktail party with different listening strategies facilitated by different types of musical ability. Unfortunately, our sample is not large enough to further stratify our listeners into instrument-specific subgroups. However, there is evidence that musicians listen and react to music differently (e.g., Mikutta et al., 2014) and show genre-specific tuning of brain activity. For example, classical musicians showing heightened P3 responses when listening to classical music, and rock musicians when listening to rock music (Caldwell & Riby, 2007). Future studies that recruit participants specifically based on primary instrument training would be needed to probe this further.

## 5. Conclusion

In summary, our results provide novel insight into how we listen to speech in background music. Listening to any music can impair concurrent speech understanding, and familiar music is particularly distracting. These differences occur as early as 50 ms during speech processing, supporting models of early-attentional control that exert influences on speech coding within the primary auditory cortices. Speech tracking is weaker when attending to background music, but only for less musical individuals. These findings reveal that exogenous properties of acoustic mixtures and endogenous factors of the listener interact when navigating noisy listening environments. Still, more research is needed to determine what aspects of musicality or listening strategies cause these differential effects.

## Acknowledgments

The authors thank Jessica MacLean and Rose Rizzi for comments on the early version of this manuscript. This work was supported by the National Institutes of Health (NIH/NIDCD R01DC016267 to G.M.B.).

